# *In-silico* identification, characterization, and expression analysis of RNA recognition motif (RRM) containing RNA binding proteins in *Aedes aegypti*

**DOI:** 10.1101/2023.07.20.549814

**Authors:** Melveettil Kishor Sumitha, Mariapillai Kalimuthu, Murali Aarthy, Rajaiah Paramasivan, Ashwani Kumar, Bhavna Gupta

**Author notes:** **Corresponding author:** Bhavna Gupta, Scientist D, ICMR-Vector Control Research Centre (VCRC), Field Station, 4, Sarojini Street Chinna Chokkikulam, Madurai-625002.

## Abstract

**Background:** RNA binding proteins (RBPs) are the proteins that bind RNAs and regulate their functioning. RBPs in mosquitoes are gaining attention due to their ability to bind flaviviruses and regulate their replication and transmission. Despite their relevance, RBPs in mosquitoes have not been explored much. In this study, we screened the whole genome of *Aedes aegypti,* the primary vector of several pathogenic viruses, and identified the proteins containing RNA recognition motif (RRM), the most abundant protein domain in eukaryotes.

**Results:** Using several *in-silico* strategies, a total of 135 RRM containing RBPs were identified in *Ae. aegypti*. The proteins were characterized based on their available annotations and the sequence similarity with the *Drosophila melanogaster*. *Ae. aegypti* RRM containing RBPs included serine/arginine-rich (SR) proteins, polyadenylate binding proteins (PABP), heteronuclear ribonucleoproteins (hnRNP), small nuclear ribonucleoproteins (snRNP), splicing factors, eukaryotic initiation factors, transformers, nucleolysins, *etc*. Phylogenetic analysis revealed that the proteins and the domain organisation is conserved among *Ae. aegypti*, *Bombyx mori* and *Drosophila melanogaster*. However, the gene length and the intron-exon organisation varied across the insect species. Expression analysis of the genes encoding RBPs using publicly available RNA sequencing data for different developmental time points of the mosquito life cycle starting from the ovary and eggs up to the adults revealed stage-specific expression with several genes preferentially expressed in early embryonic stages and blood-fed female ovary.

**Conclusions:** This is the first database for the *Ae. aegypti* RBPs that can serve as the reference base for future investigations. The stage-specific genes can be further explored to determine their role in mosquito growth and development with a focus to develop novel mosquito control strategies.

## Introduction

RNA binding proteins (RBPs) are the proteins that bind RNAs and regulate their life cycle starting from synthesis to decay. RNAs after transcription are always associated with RBPs and RBPs along with other cis-regulatory elements regulate RNA processing, transportation, localization, stability, modifications, translation, and degradation (Glisovic et al. 2008). Due to their role in several aspects of RNA biology, Diaz-Munoz and Turner in 2018 have aptly described RBPs as ‘readers, writers, editors, and erasers of the transcriptome’ where ‘RNA writers’ include the proteins involved in splicing, capping, and polyadenylation, ‘readers’ are the RBPs involved in subcellular localization and translation, ‘editors’- constitute methyltransferases and deaminases, and ‘erasers’- include the destabilizing factors and nucleases that cause RNA instability and degradation (Díaz-Muñoz and Turner 2018). The regulation of all these aspects after transcription is known as post-transcriptional gene regulation (PTGR) and RBPs play a major role in PTGR.

RBPs constitute a significant part of the proteome in eukaryotes and are known to regulate cell differentiation and homeostasis in response to different developmental stages, environmental challenges, and diseases/infections (Wurth 2012; Brinegar and Cooper 2016; Li et al. 2020). Several RBPs have been associated with different types of cancers, and genetic disorders (Wurth 2012; Li et al. 2020). RBPs also play an important role in the development and growth processes of organisms as seen in plants (Muleya and Marondedze 2020), animals (Kerner et al. 2011), bacteria (Holmqvist and Vogel 2018), and insects (Norvell et al. 1999; Yoon et al. 2018). Interestingly, many RBPs have been found to regulate early embryo development in *Drosophila*. For example, mutations in the *Squid* gene disrupted Gurken-dependent dorsoventral patterning during oogenesis in *Drosophila* resulting in female sterility (Norvell et al. 1999; Gamberi et al. 2006). Smaug, another RBP has been found to be involved in maternal to zygotic transition (Benoit et al. 2009). The localization and translational regulation of maternal transcripts during oogenesis in *Drosophila* relies on RBPs (Bansal et al. 2020). Staufen, a double-stranded RBP, has been identified as a key player in RNAi in certain Coleopteran insects (Yoon et al. 2018). However, RBPs in mosquitoes and their possible functions have not been explored much except the reports where mosquito RBPs have been found to control the replication of flaviviruses (Diosa-Toro et al. 2020; Yeh et al. 2022). RNA viruses use host RBPs for their replication and survival and such RBPs can act as antivirals. For example, AeStaufen reduced genomic and sub-genomic flaviviral RNA copies in several mosquito tissues, including the salivary glands indicating AeStaufen’s role in dengue transmission (Yeh et al., 2022). Taschuk et al.,(2020) described the antiviral role of DDX56, Dead box helicase in *Drosophila* and human cell lines (Taschuk et al. 2020). Thus, identification and characterization of mosquito RBPome would be helpful for such studies.

RBPs are generally identified by the presence of different structural motifs which locate and interact with their specific RNA targets. RNA recognition motif (RRM) is one of the most abundant and well-characterized RNA binding motif in eukaryotes (Venter et al. 2001). RRM has been found in all forms of life including bacteria and viruses. RRM containing proteins are known to be involved in several post-transcriptional aspects like alternative splicing, translation, degradation, and processing of precursor ribosomal RNA (pre-rRNA). RRM containing proteins form a prominent class of RBPs performing important biological functions.

In this study, we screened *Ae. aegypti* genome and identified the RRM containing proteins using various *in-silico* approaches. We also carried out expression analysis of the genes encoding the RBPs to elucidate their association with mosquito growth and developmental stages using stage-specific transcriptome data (Akbari et al. 2013).

## Material and Methods

### Identification and characterization of RRM-containing proteins in *Ae. aegypti*

To search for RRM-containing proteins in *Ae. aegypti* genome, we used multiple *in-silico* approaches. First, ‘RRM and *Ae. aegypti*’ was used as the keyword in GenBank to retrieve proteins containing RRM. Second, 134 RRM containing proteins of *D. melanogaster* (Gamberi et al. 2006) were used as a protein query against *Ae. aegypti* in BLASTp search. Third, the UniProt entries for *Ae. aegypti* for RRM-containing proteins were retrieved. Fourth, RRM profiles from PROSITE for IDs PDOC00030, PDOC51472, and PDOC51939 were used as a query for BLASTp against *Ae. aegypti* database with default parameters. The proteins obtained from each strategy were carefully examined to remove the duplicates and lists of unique proteins were obtained. This list was further used in the BLASTp search against *Ae. aegypti* protein database to retrieve proteins that might have been missed from earlier searches.

The list of proteins finally obtained was analyzed using SMART and Conserved Domain (CD) databases to confirm the presence of RRM and the domain organisation for each protein was retrieved from the SMART database. The chromosomal locations, protein length, number of transcripts and protein IDs were obtained from the VectorBase (https://vectorbase.org/). Functional annotation was done using Blast2GO (Conesa et al. 2005) and the descriptions for each protein were based on available annotations in Genbank/Vectorbase or sequence similarity with *D. melanogaster* (>30% identity).

### Phylogenetic analysis, chromosomal mapping, and gene structure

Phylogenetic analysis of full-length amino acid sequences of *Ae. aegypti* RBPs were aligned using CLUSTAL W and Neighbour-joining tree with 500 bootstraps was constructed using MEGA7.0 (Kumar et al. 2016). The most similar proteins in *D. melanogaster* and *Bombyx mori* were retrieved from VectorBase (https://vectorbase.org/vectorbase/app) and Ensembl (https://metazoa.ensembl.org/index.html), respectively. Annotations were added to the tree using the iTOL server (Letunic and Bork 2021). Chromosomal mapping of the genes on *Ae. aegypti* chromosomes were done using Mapchart 2.32 (Voorrips 2002). Intron-exon gene structure schematics were generated using GSDS an online server (Hu et al. 2015).

### Expression profiling of the genes encoding RRM containing proteins

RNA sequencing data from 42 different developmental time points throughout the *Ae. aegypti* life cycle from eggs up to the adult stage was obtained from Sequence Read Archive (SRA) a public repository database, with accession number SRP026319 (Akbari et al. 2013). Transcriptomic data were mapped to the *Ae. aegypti* AaegL5.0 reference genome using HISAT2 (Kim et al. 2019) to identify their genomic positions. The reads mapped to each gene were assembled and quantified using Cufflinks (Trapnell et al. 2012). The read count was quantified as the number of fragments per kilobase of exon per million mapped fragments (FPKM). The FPKM for *Ae. aegypti* RRM containing RBPs were used for clustering and heatmap generation using Morpheus, (https://software.broadinstitute.org/morpheus).

## Results and Discussion

### Overview of RBPs in *Ae. aegypti*

Using multiple *in-silico* strategies, *Ae. aegypti* genome was screened and a total of 135 RRM containing RBPs were identified (Supplementary Table 1) which is comparable to the number of RBPs in *D. melanogaster* (n=134) (Gamberi et al. 2006; Sysoev et al. 2016) and *B. mori* (n=123) (Wang and Zhou 2009). Orthologs of 133 RBPs were identified in *D. melanogaster* (Supplementary Table 1). Annotations were taken as available in Genbank or VectorBase and the remaining unannotated genes were putatively characterized based on sequence similarity with *D. melanogaster*. A total of 126 proteins could be described (Supplementary Table 1), while 9 RBPs could not be assigned any description. 6 of these 9 RBPs contained multiple RRMs, such as AAEL022693 and AAEL012243 had 6 RRM copies; AAEL019864 and AAEL023907 contained 5 copies; AAEL022113 and AAEL004699 had 4 copies. 3 of the 9 RBPs contained single RRM (AAEL022876, AAEL025927, AAEL019879).

RRM is often found as a single copy or in multiple copies and is also commonly found in combination with other motifs (Maris et al. 2005; SenGupta 2013; Loerch and Kielkopf 2015). Out of 135 RBPs, 41 (30%) had a single RRM and 52 proteins had multiple RRMs (37%). The remaining 42 RBPs contained RRM along with other motifs (33%). Zinc finger (Znf) was the most common as earlier seen in many other organisms (Maris et al. 2005; Mahalingam and Walling 2020). Znf is a small protein motif with multiple finger-like protrusions that make tandem contact with their target molecule. There are many superfamilies of Znf that have binding affinities to DNA/RNA/proteins based on their sequence and structure. For example; CCHHs is known to bind double-stranded DNA/RNA, and CCCH and CCHC types bind single-stranded RNA (Summers 1991; Iuchi 2001; Michel et al. 2003; Wang et al. 2021). A total of 13 Znf motifs including five C3H1, three C2HC, two C2H2, one ring finger, and three RBZ were observed in 11 proteins. The ring finger is involved in protein-protein interaction and the Znf-RBZ is related to the ubiquitin-binding function (Lorick et al. 1999).

Other motifs included PWI, KH, and G-patch, which are also known for their RNA binding affinities (Szymczyna et al. 2003; Dong et al. 2004; Valverde et al. 2008; Aksaas et al. 2011; Zhang et al. 2016). Apart from this, motifs like SURP, SAP, LA, SPOC, RPR, HAT, Lsm_interact, methyltransferase, muHD, CID are known to be involved in transcriptional and translational processes. For example; SURP is found in splicing regulatory proteins (Kuwasako et al. 2006). SAP motif has been identified in the proteins involved in DNA repair, RNA processing, and apoptotic chromatin degradation (Aravind and Koonin 2000). Three proteins contained the LA motif which is known to be involved in the maturation of RNA polymerase III transcripts (Alfano et al. 2004; Dong et al. 2004). LA-motif also recognizes mRNAs with 5′-terminal oligopyrimidine motif (5’TOP) important for protein synthesis (Pellizzoni et al. 1997). One protein had RRM in combination with the cleavage stimulation factor (CSTF) which is known for 3’ end cleavage and polyadenylation of pre-mRNAs (Mandel et al. 2008). HAT, a helical repeat motif, is involved in RNA binding and RNA remodelling (Hammani et al. 2012). Nuclear transport-like factor (NTF2) was identified in one *Ae. aegypti* RBP. NTF2 functions as a cytosolic factor for nuclear import and also interacts with nuclear pore complex protein (Paschal and Gerace 1995). The UBA (ubiquitin-binding associated) was found in one of the proteins and the UBA mediates ubiquitination (Hurley et al. 2006). Presence of different motifs having specific functions modulates the binding affinity, specificity, and versatility of RBPs. They may also be responsible for locating and interacting with multiple RBPs.

RRMs are also commonly seen in multiple copies within a protein. 52 proteins contained multiple RRMs ranging from two to seven. Among proteins with multiple RRM copies, AAEL021950 contained the maximum number of seven RRMs, and four RBPs namely, AAEL004075, AAEL012243, AAEL020196, and AAEL022693 contained six RRMs. Some of the RBPs are known to have a specific number of copies. For example, a single RRM is found in spliceosomal U1 70 kDa protein, two in U2 auxiliary factor 65 kDa subunit, three RRM copies are generally seen in ELAV protein, and the polyadenylate binding proteins (PABP) are known to have four RRM copies (SenGupta 2013). Although the function of each RRM among multiple copies is unknown; they may increase specificity by binding to the long stretch of RNA sequence and function cooperatively (Maris et al. 2005) or may have different specificities and thus diversify the biological functions of the protein. Moreover, it is also possible that all of the RRMs may not be functional or may not be able to bind RNAs (SenGupta 2013).

### Chromosomal mapping and functional enrichment

RBPs were mapped onto the *Ae. aegypti* chromosomes and 135 RBP encoding genes were found evenly distributed across the length of all the 3 chromosomes (Figure 1). The largest Chromosome (chr_2) had a maximum of 53 genes, chromosome 3 had 45 and the smallest chromosome (chr_1) contained 37, the lowest number of genes. The length of proteins varied from 91 to 5680 amino acids including 43 small proteins (<320 amino acids) and 32 large proteins (>700 amino acids). The majority (75%) of the proteins that had only one exon were small proteins. The number of predicted transcripts for each protein varied from 1 to 29 and 56% (76/135) of the total genes had only one transcript.

**Figure 1.**
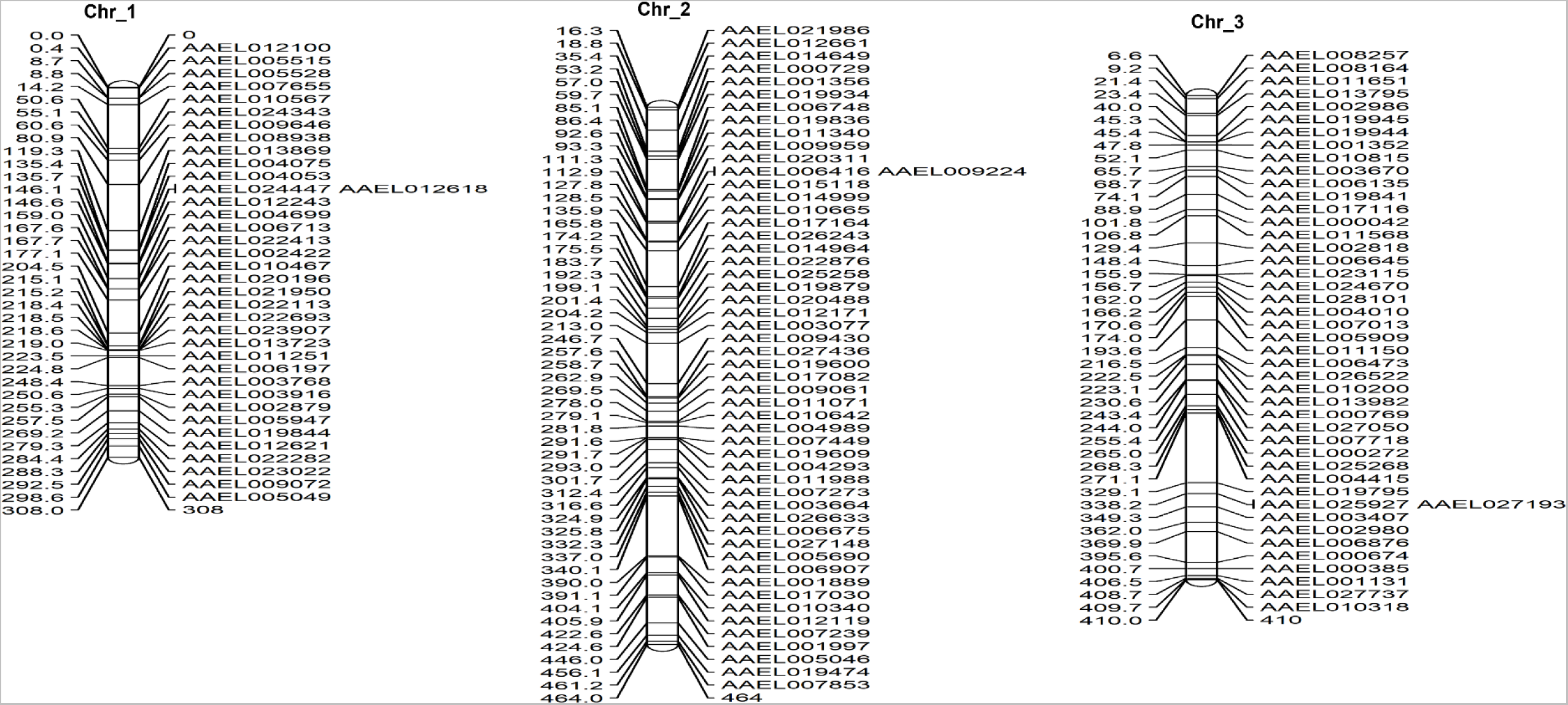
Chromosomal distribution of the genes encoding RNA binding proteins containing RNA recognition motif (RRM) across three chromosomes (chr_1, chr_2 & chr_3) of *Aedes aegypti*. The numbers on the left of each chromosome are the locations of the genes (in Mb) and the protein IDs are shown on the right side.

Gene Ontology (GO) analysis with BLAST2GO categorized RBPs based on their putative functions. As expected, ‘RNA binding’ was the most enriched term among molecular functions, and in biological processes, *Ae. Aegypti* RBPs were found associated with ‘gene expression regulation’, ‘nucleic acid and protein metabolism’, ‘macromolecule biosynthesis’ ‘cellular component assembly’, ‘protein containing complex’, macromolecule modification’, and ‘ribonucleoprotein complex’. This indicates the involvement of RBPs in several biological processes as seen in several other organisms (Figure 2).

**Figure 2.**
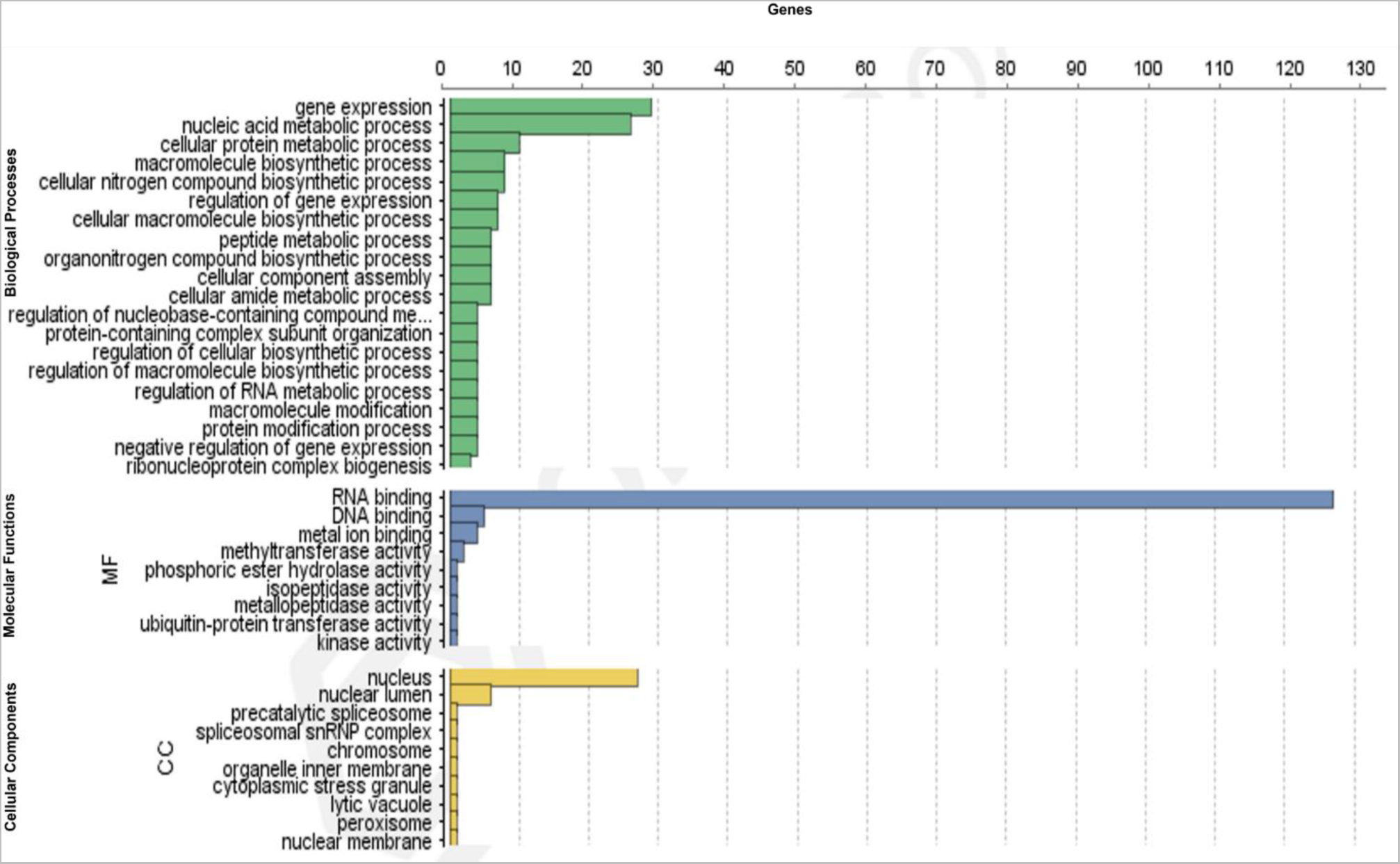
Gene ontology classifications of the biological process, molecular function, and cellular localization of the genes encoding RNA binding proteins containing RNA recognition motif (RRM) in *Aedes aegypti*. GO distribution for the top 20 levels has been shown.

### Expression analysis of RBP genes using RNA sequencing data

Since developmental stage-specific transcriptomes of *Ae. aegypti* were available in a public database for 42 time points throughout its life cycle starting from early embryo up to adult mosquitoes (Akbari et al. 2013), it was possible to explore the stage-specific expression of RBPs in *Ae. aegypti*. From the analysis of RNA seq data, we retrieved FPKM values for 131 RBP encoding genes, and all the genes were found to have >1.0 FPKM in more than one stage and thus were considered expressed. Hierarchical clustering of the gene expression data indicated that the majority of the genes exhibited preferential expression in the embryonic stages and blood-fed female ovaries (Figure 3). Only one gene (AAEL007013: RNA binding protein precursor) was found expressed in all the stages. A subset of RBPs was found highly expressed in early embryonic stages and interestingly those RBPs were also found highly expressed in blood-fed ovaries (Figure 3). This indicates that these RBPs might be maternally inherited and possibly plays essential roles in embryonic development which is in corroboration with the findings that RBPs are essentially involved in oogenesis and embryo development. The *Ae. aegypti* eggshell proteome analysis by Marinotti et al, (Marinotti et al. 2014) also identified two RBPs (AAEL010665 and AAEL013869) suggesting their possible role in egg development. The detailed characterization of these genes is warranted to identify the candidates for developing new mosquito growth regulators or insecticides.

**Figure 3.**
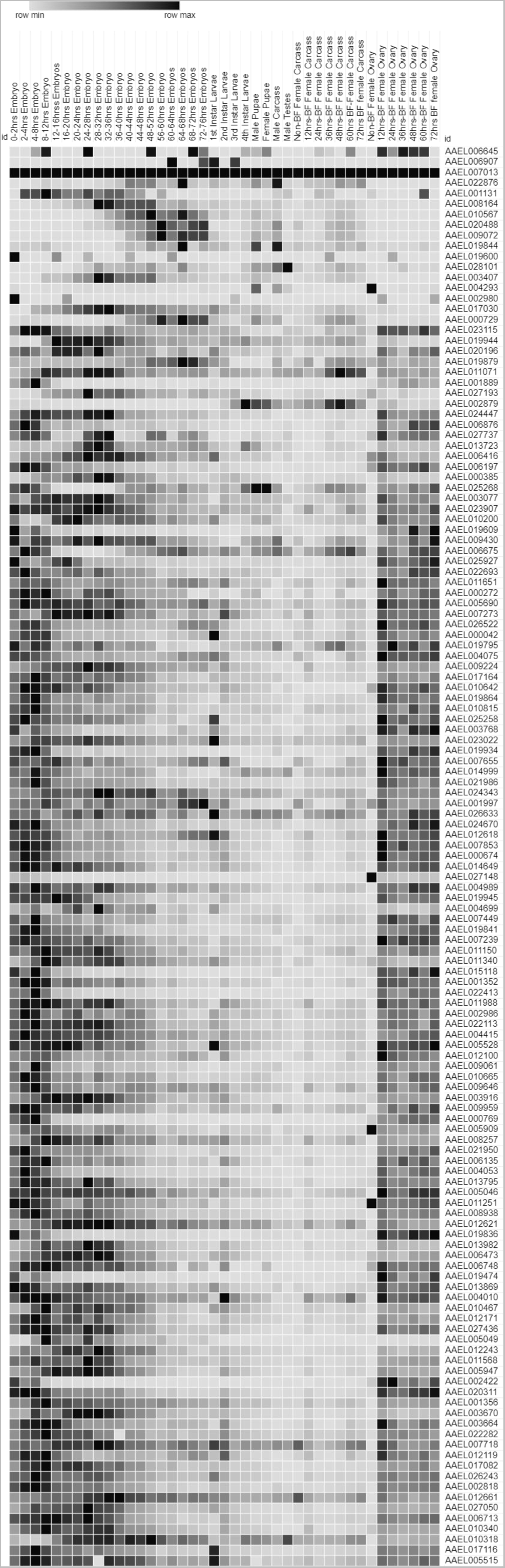
Expression profiles of the genes encoding RNA binding proteins containing RNA recognition motif (RRM) in *Aedes aegypti* across 41 developmental time points starting from eggs up to adult mosquitoes. These developmental stages include specific time intervals of embryonic development, larval stages, sex-biased expression of male versus female pupae and carcass, blood-fed versus non-blood-fed ovary and female carcass, and testes-specific expression. Colors from white to black represent increasing gene expression

### Phylogenetic relationships, gene structure and domain organisation

RRM-containing proteins belonged to a wide range of functional categories including; serine arginine (SR) rich proteins, heterogeneous nuclear RNP (hnRNPs), small nuclear ribonucleoproteins (snRNPs), nucleolysins, polyadenylate binding proteins (PABP), ELAV like proteins, eukaryotic translation initiation factors, etc. (Table 1). To comprehensively analyse evolutionary relationships, we analysed gene structure, domain organisation and phylogenetics of some of the major *Ae. aegypti* RBP classes and in comparison, with *D. melanogaster* and *B. mori*.

**Table 1.** Summary of the functional classes of proteins and the number of genes in each class.

| **Sl. No.** | **Functional class** | **Number of genes** |
| --- | --- | --- |
| 1 | Serine/Arginine rich proteins | 12 |
| 2 | Zinc finger proteins | 6 |
| 3 | Small nuclear ribonuclear proteins | 8 |
| 4 | Pre-mRNA splicing factors | 3 |
| 5 | Heterogenous nuclear ribonuclear proteins | 8 |
| 6 | Poly(A)-binding protein | 5 |
| 7 | Cytoplasmic polyadenylation element binding proteins (cpeb) | 2 |
| 8 | ELAV like proteins | 5 |
| 9 | Eukaryotic translation initiation factor | 4 |
| 10 | Nucleolysin | 1 |
| 11 | Peptidyl-propyl cis-trans isomerase | 2 |
| 12 | Musashi like proteins | 2 |
| 13 | Splicing proteins | 5 |
| 14 | Transformer proteins | 4 |
| 15 | La proteins | 3 |
| 16 | SRA stem-loop interacting RBPs | 3 |
| 17 | Sex lethal | 1 |
| 18 | Other characterised RBPs | 53 |
| 19 | Unknown RBPs | 8 |

### Spliceosome-associated proteins

pre-mRNA splicing needs core spliceosomes, cis-sequence elements, and several RBPs (Fredericks et al. 2015; Vuong et al. 2016). Among RBPs, snRNPs form the core components of the spliceosomes, and SR and hnRNPs proteins are the major splicing factors. A total of 36 such proteins with a putative role in splicing were identified including 12 SR proteins, eight hnRNPs, three pre-mRNA splicing factors, eight snRNPs, and five other splicing factors (Supplementary Table1). Brooks and co-workers (Brooks et al. 2015) reported 56 *Drosophila* RBPs involved in splicing. These 56 proteins were BLAST searched against *Ae. aegypti* genome to identify the similar proteins. and identified total 37 proteins containing RRM. These 37 proteins belonged to different categories including six SR proteins, two pre-mRNA splicing factors, one each from hnRNP, PABP, ELAV along with two nucleolysins, two translation initiation factors, etc. This is possible as many RBPs are known to perform multiple functions, however, the functional characterization of *Ae. aegypti* may provide accurate information.

### Serine arginine-rich (SR) proteins

A total of 12 SR proteins were identified in *Ae. aegypti*. SR proteins contain SR repeat regions at the C terminal and RRM at the N terminal. Nine proteins had a single RRM and five had two RRMs. One of the proteins contained Znf along with RRM and another protein had an RPR motif which helps in protein-protein interaction. SR proteins are known to be a part of spliceosome assembly and regulate the alternate splicing (Jeong 2017) while RRM in the protein binds to the splicing enhancers to regulate processing events and the SR rich region binds to the other proteins or RNAs.

We analysed the 12 *Ae. aegypti* putative SR proteins along with 10 SR proteins from *D. melanogaster* and 9 from *B. mori*. Phylogenetic analysis revealed that SR proteins are highly conserved among the three insect species. *Ae. aegypti* proteins clustered closely with their orthologs from both *D. melanogaster* and *B. mori* (Figure 4). Protein length and the domain organisation were found conserved (Matthews et al. 2018), while the gene length and the intron-exon structure were highly variable across the species. Number of introns and their organisation also varied among the species. These findings are in corroboration with the findings of genomic comparisons between *Ae. aegypti, Anopheles gambiae and D. melanogaster* (Nene et al. 2007) which observed that *Ae. aegypti* genes are generally longer and contain longer introns. This suggested that the genes have evolved across the species but they are highly conserved at protein level indicating the biological importance of these proteins and the conserved functional role played by them.

**Figure 4.**
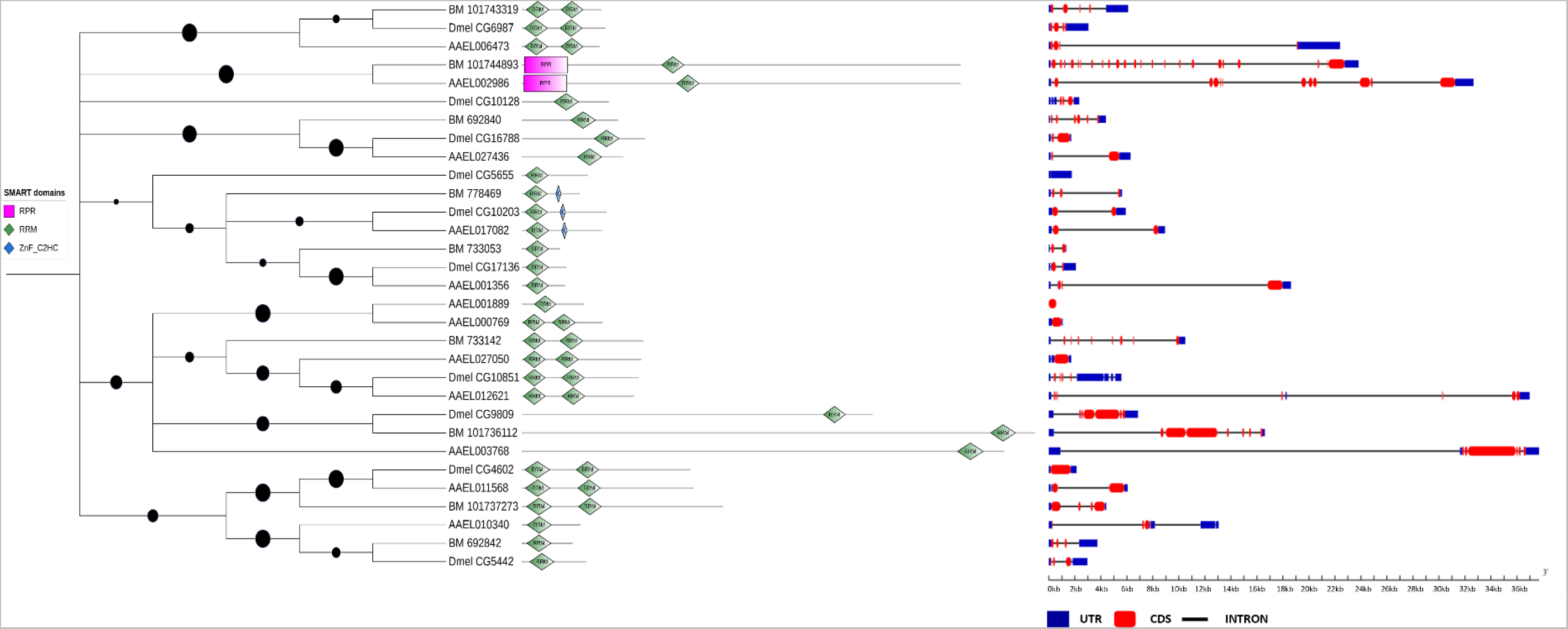
Phylogenetic tree of *Aedes aegypti, Drosophila melanogaster and Bombyx mori* Serine/Arginine rich proteins containing RNA recognition motif (RRM). The tree was constructed using the Neighbour Joining (NJ) method with 500 bootstraps. The domain architecture and the intron-exon architecture of each protein is also shown. Bootstrap values are represented by the size of the red circle on each branch.

### Small nuclear ribonucleoproteins (snRNPs)

Removal of introns from pre-mRNA (splicing) is a crucial step in eukaryotes’ gene expression. This is carried out by a ribonucleoprotein complex called spliceosome which is comprised of snRNPs and various other proteins. Seven snRNPs containing the RRM were identified in *Ae. aegypti*. snRNP is a complex of proteins and snRNA. snRNPs associated with specific U-rich snRNAs are the core components of the spliceosomes involved in splicing (Solymosy and Pollák 1993). The most abundant and well-characterized snRNPs include; U1, U2, U4, U5, and U6 which form the major components of spliceosomes (Solymosy and Pollák 1993) and the minor spliceosomes contain U11, U12, and U4atc, U5 and U6atac (Will and Lührmann 2011; Matera and Wang 2014). snRNP in *Ae. aegypti* included 3 U1 snRNPs, one U11/12, two proteins with U2 auxiliary factor and one with U2 associated SURP motif containing protein. Phylogenetic analysis of these snRNPs along with *D. melanogaster* and *B. mori* revealed the similar pattern as observed in SR proteins. The proteins and the domain organisation were highly conserved but the variations were observed in gene length and intron-exon organisation (Figure 5).

**Figure 5.**
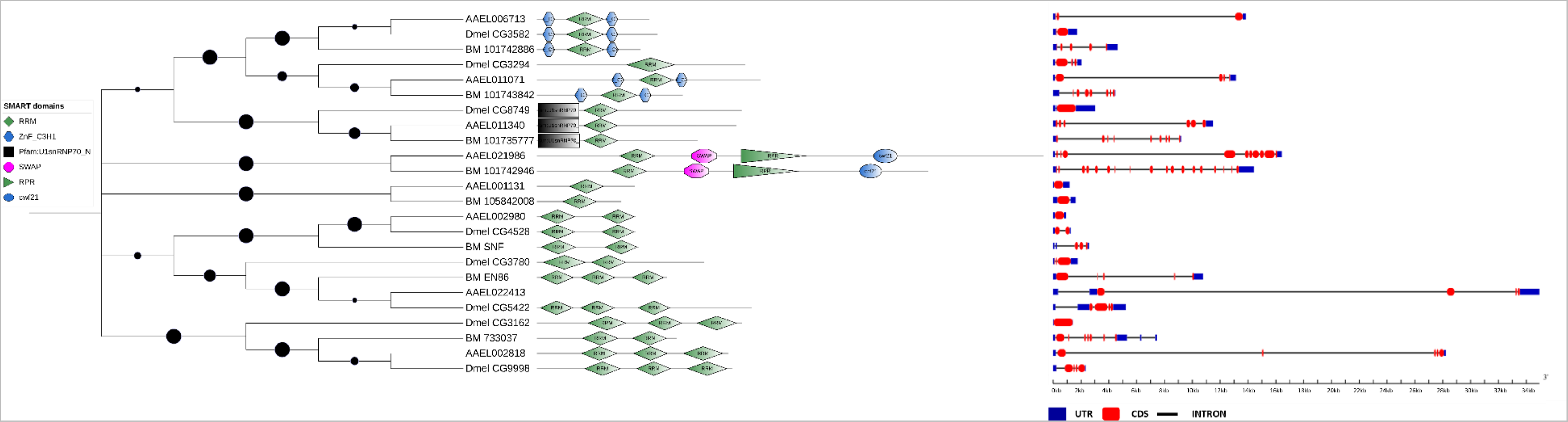
Phylogenetic tree of *Aedes aegypti, Drosophila melanogaster and Bombyx mori* small nuclear RNA nucleoproteins (snRNPs) containing RNA recognition motif (RRM). The tree was constructed using the Neighbour Joining (NJ) method with 500 bootstraps. The domain architecture and the intron-exon architecture of each protein is also shown. Bootstrap values are represented by the size of the red circle on each branch.

### Heterogeneous nuclear ribonucleoproteins (hnRNPs)

The RRM is also found in several hnRNPs. The majority of hnRNPs are known to regulate alternative splicing and protein components of snRNPs. hnRNPs associate with nascent RNAs and lead to their export, translation, localization, and stability (Krecic and Swanson 1999; Geuens et al. 2016). A total of eight hnRNPs were identified, five of which encoded two RRMs and three proteins had three RRMs. All the proteins clustered in a phylogenetic tree as per their domain organisation (Figure 6). Two of the *Ae. aegypti* hnRNPs (AAEL005515 and AAEL005049) were clustered with *Drosophila* squid protein. Squid is the most abundant hnRNP protein which is expected to bind most cellular RNAs. *Squid* plays an important role in dorso- ventral axis formation during oogenesis by localizing *Gurken*RNA (Norvell et al. 1999; Steinhauer and Kalderon 2005; Cáceres and Nilson 2009). Several mutants of this gene have been generated and have shown the role of the gene in important aspects of oogenesis and embryo development. For example, homozygous mutants laid severely dorsal eggs that could not mature into adulthood (Kelley 1993; Matunis et al. 1994), complete deletion of *squid* genes lead to lethality (Matunis et al. 1994), while deletion in the first exon of the *squid* resulted in larval death (Matunis et al. 1994). *Ae. aegypti* proteins AAEL005515 and AAEL005049 have not been characterized. The amino acid sequences of both the proteins >75% identical to the *Drosophila* squid protein (Supplementary Figure 1). However, gene structure and the overall gene length was different (Figure 6). Protein-protein interaction analysis with STRING database (Szklarczyk et al. 2021) using default parameters revealed interactions between AAEL005515 and AAEL005049 as well as with the other nine proteins (Figure 7). Seven of the nine proteins coded for RNA-binding proteins. Particularly, AAEL005515 and AAEL005049 were found interacting with all other nine genes indicating their importance in the network of proteins. The sequence conservation across the species indicates the functional relevance, however, in-depth analysis of these genes could provide greater insights and envision.

**Figure 6.**
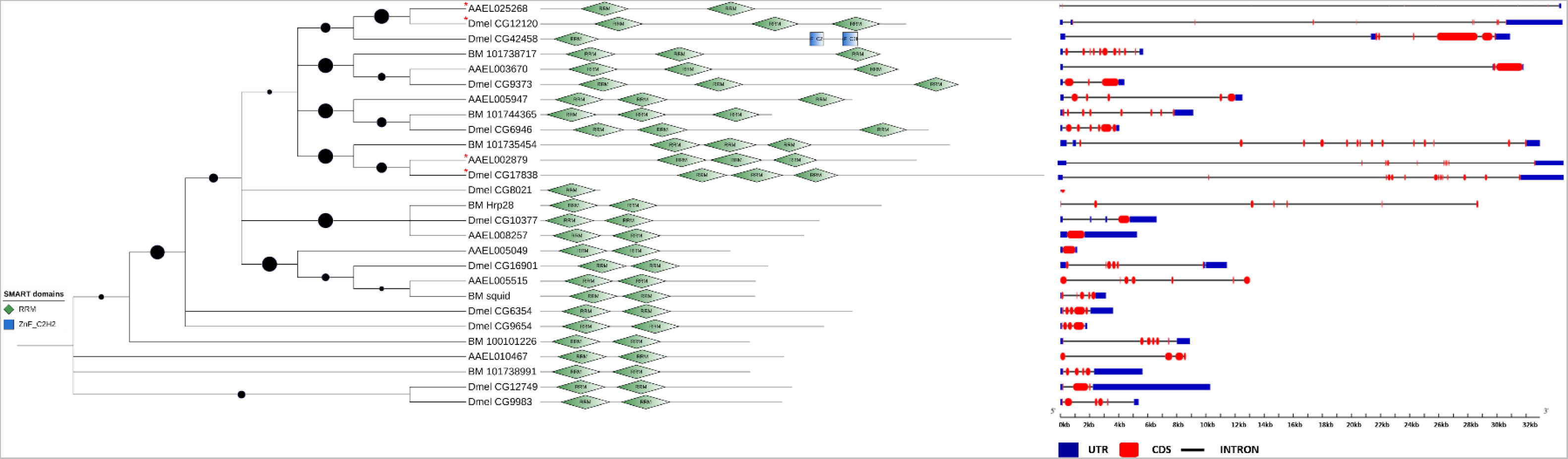
Phylogenetic tree of *Aedes aegypti, Drosophila melanogaster and Bombyx mori* heterogeneous nuclear ribonucleoproteins (hnRNPs) containing RNA recognition motif (RRM). The tree was constructed using the Neighbour Joining (NJ) method with 500 bootstraps. The domain architecture and the intron-exon architecture of each protein is also shown. Bootstrap values are represented by the size of the red circle on each branch.

**Figure 7.**
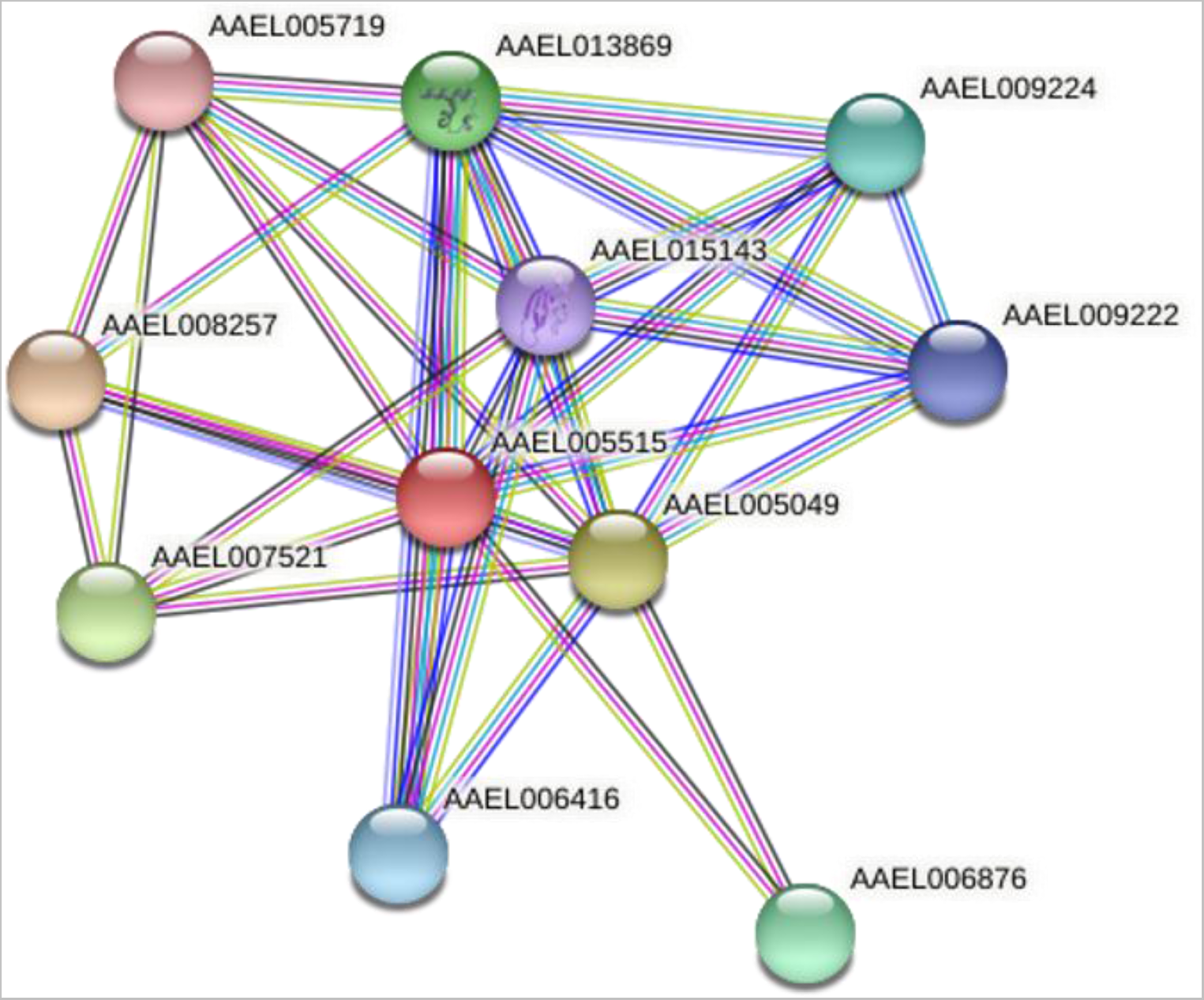
The STRING protein-protein interaction network for *Aedes aegypti* protein AAEL005515. Colored lines between the proteins indicate interactions.

### Polyadenylate binding proteins (PABP)

PABPs, another RRM containing RBPs are involved in protein synthesis, mRNA stability, and mRNA biogenesis (Bernstein et al. 1989; Kühn and Wahle 2004). As per published records, only one PABP has been predicted in *D. melanogaster* (Smith et al. 2014), three in humans, while 12 in *Arabidopsis thaliana* (Lorković and Barta 2002), five in rice (Belostotsky 2003) and 14 in barley (Mahalingam and Walling 2020). In *Ae. aegypti*, we identified five PABP and two cytoplasmic polyadenylation element binding (CPEB) proteins. Similar to other eukaryotes, four PABPs had multiple RRMs ranging from two to six. However, only one of them contained a polyA domain with 4RRMs which is the well characterized domain pair identified in PABPs. One protein had 2RRMs and three had single RRM copy. Similar to plants, where PABPs with one or 2RRMs have been identified (Lorković and Barta 2002; Mahalingam and Walling 2020), however, the functional characterization of the proteins could shed light on their exact role and functional capabilities.

## Conclusion

This study has generated the first reference database for the *Ae. aegypti* RBPs containing RRMs and their expression analysis specific to different developmental time points of the mosquito life cycle. This database would be useful for future studies to select and characterize the candidate proteins and elucidate their role in mosquito growth and development. The proteins having a significant impact on the growth and development of mosquitoes can be targeted to develop mosquito growth regulators or insecticides. This will also be a useful reference base for future studies to characterize the entire RBP repertoire of the *Ae. aegypti* as well as other vector species. It can further be extended to identify and validate the genes experimentally through interactome capture from the different developmental stages of mosquitoes to identify candidate genes.

## Acknowledgments

Authors would like to thank the Indian Council of Medical Research (ICMR), New Delhi for intramural support. Melveettil Kishor Sumitha would like to thank ICMR for the Senior Research Fellowship and Madurai Kamaraj University (MKU) for supporting the research.

## Availability of data and material

All the data has been included in the manuscript.

## Author Contributions

Conceptualization, Methodology, Formal analysis and investigation, writing manuscript: Bhavna Gupta, Resources, manuscript review: Ashwani Kumar and Rajaiah Paramasivan, Methodology and Data Curation: Melveettil Kishor Sumitha, Mariapillai Kalimuthu, and Murali Aarthy

## Ethics approval

Not Applicable

## Consent to participate

Not Applicable

## Consent for Publication

Not Applicable

