## Supplementary files for "*In-silico* identification, characterization, and expression analysis of RNA recognition motif (RRM) containing RNA binding proteins in *Aedes aegypti*"

***Corresponding author:**

**Supplementary Table 1. Summary of RRM containing RNA binding proteins in _Aedes aegypti_**

| Sl. No. | Gene name | Domain | Length | Transcripts | Product Description | Most similar protein in _D. melanogaster_ |
|---|---|---|---|---|---|---|
| **Serine/Arginine rich proteins** | | | | | | |
| 1 | AAEL012621 | 2RRM | 140 | 4 | serine-arginine protein 55 | Dmel_CG10851, B52 (SR55) |
| 2 | AAEL006473 | 2RRM | 237 | 2 | arginine/serine-rich splicing factor 1A | Dmel_CG6987, SF2 |
| 3 | AAEL000769 | 2RRM | 245 | 1 | arginine/serine protein 55 | Dmel_CG10851, SR55 |
| 4 | AAEL010340 | RRM | 177 | 3 | serine/arginine-rich splicing factor 2 | Dmel_CG5442, SC35, |
| 5 | AAEL001356 | RRM | 131 | 3 | RNA-binding protein 1 | Dmel_CG17136, RBP1 |

| | | | | | | |
|---|---|---|---|---|---|---|
| 6 | AAEL017082 | RRM, ZnF_C2HC | 243 | 1 | serine/arginine-rich splicing factor 6 | Dmel_CG10203, x16 |
| 7 | AAEL027050 | 2RRM | 363 | 1 | serine-arginine protein 55-like | Dmel_CG10851, SR55 |
| 8 | AAEL027436 | RRM | 308 | 1 | RNA binding protein with serine-rich domain 1-A-like | Dmel_CG16788, RnpS1 |
| 9 | AAEL001889 | RRM | 189 | 1 | serine-arginine protein 55 | Dmel_CG10851, SR55 |
| 10 | AAEL002986 | RPR, RRM | 1334 | 3 | protein SCAF8 | Dmel_CG4266 |
| 11 | AAEL003768 | RRM | 1467 | 4 | serine/arginine repetitive matrix protein 2 | Dmel_CG9809, spargel |
| 12 | AAEL011568 | 2RRM | 522 | 1 | arginine/serine-rich7, probable splicing factor | Dmel_CG4602, Srp54 |
| **Zinc finger proteins** | | | | | | |
| 13 | AAEL006197 | RRM, ZnF_C2HC | 207 | 1 | zinc finger CCHC-type and RNA-binding motif-containing protein 1 | Dmel_CG8597, Lark |
| 14 | AAEL009061 | PWI, RRM, Nup35_RRM_2 | 1103 | 2 | zinc finger protein swm | Dmel_CG10084, swm |
| 15 | AAEL019600 | RRM, 2ZnF_RBZ | 389 | 1 | RNA-binding protein cabeza | Dmel_CG3606, Cabeza |
| 16 | AAEL022282 | RRM, ZnF_RBZ | 395 | 5 | RNA-binding protein Cabeza-like | Dmel_CG3606, Cabeza |
| 17 | AAEL013982 | 2RRM, ZnF_C2HC | 330 | 1 | RNA binding motif protein, lark | Dmel_CG8597, lark |

| | | | | | | |
|---|---|---|---|---|---|---|
| 18 | AAEL004989 | 2RRM, ZnF_RBZ, ZnF_C2H2, G Patch | 907 | 1 | RNA-binding protein 5-A | Dmel_CG9373, rumpelstiltskin |
| **Small nuclear ribonucleoproteins** | | | | | | |
| 19 | AAEL011340 | U1snRNP70_N, RRM | 437 | 1 | U1 small nuclear ribonucleoprotein 70 kDa | Dmel_CG8749, snRNP-U1-70K |
| 20 | AAEL002980 | 2RRM | 216 | 1 | U1 small nuclear ribonucleoprotein A | Dmel_CG4528, snf |
| 21 | AAEL001131 | RRM | 214 | 1 | U11/U12 small nuclear ribonucleoprotein 35 kDa protein-like | Dmel_CG8597, Lark |
| 22 | AAEL006713 | 2ZnF_C3H1, RRM | 246 | 1 | splicing factor U2af 38 kDa subunit | Dmel_CG3582, U2af38 |
| 23 | AAEL011071 | 2ZnF_C3H1, RRM | 315 | 2 | U2 snRNP auxiliary factor 35kDa subunit –related protein 2 | Dmel_CG3294, CG3294 |
| 24 | AAEL021986 | Surp, CID, RRM_1 | 1108 | 1 | U2 snRNP-associated SURP motif-containing protein-like | Dmel_CG9346 |
| 25 | AAEL022413 | 3RRM | 450 | 1 | nucleolysin TIAR | Dmel_CG5422, Rox |
| 26 | AAEL002818 | 3RRM | 418 | 1 | splicing factor u2af 50kDa subunit | Dmel_CG9998, U2af50 |
| **Pre-mRNA splicing factors** | | | | | | |

| | | | | | | |
|---|---|---|---|---|---|---|
| 27 | AAEL009959 | PRO8N, PROCON, RRM_4, U5_2-snRNA_bdg, U6-snRNA_bdg, PRP8_domainIV, JAB_MPN | 2383 | 1 | pre-mRNA splicing factor prp8 | Dmel_CG8877, pre-mRNA processing factor 8 |
| 28 | AAEL011251 | ZnF_C3H1, RRM | 429 | 1 | pre-mRNA splicing factor RBM22 | Dmel_CG14641 |
| 29 | AAEL007655 | RRM | 248 | 1 | activator of basal transcription 1 | Dmel_CG32708 |
| **Heterogeneous nuclear ribonucleoprotein** | | | | | | |
| 30 | AAEL010467 | 2RRM | 362 | 3 | heterogeneous nuclear ribonucleoprotein 87F | Dmel_CG12749, Heterogenous nuclear ribonucleoprotein at 87F – Hrb87F |
| 31 | AAEL005049 | 2RRM | 287 | 1 | RNA binding protein, squid | Dmel_CG16901, squid |
| 32 | AAEL005515 | 2RRM | 325 | 8 | RNA-binding protein squid | Dmel_CG16901, squid |
| 33 | AAEL008257 | 2RRM | 398 | 1 | heterogeneous nuclear ribonucleoprotein 27c | Dmel_CG10377, heterogeneous nuclear ribonucleoprotein at 27C |
| 34 | AAEL005947 | 3RRM | 491 | 2 | heterogeneous nuclear ribonucleoprotein H | Dmel_CG6946, gloround |
| 35 | AAEL025268 | 2RRM | 522 | 11 | heterogeneous nuclear ribonucleoprotein L | Dmel_CG9218, smooth |
| 36 | AAEL002879 | 3RRM | 561 | 29 | heterogeneous nuclear ribonucleoprotein R | Dmel_CG17838, syncrip |

| | | | | | | |
|---|---|---|---|---|---|---|
| 37 | AAEL003670 | 3RRM | 541 | 2 | heterogeneous nuclear ribonucleoprotein M | Dmel_CG9373, rumpelstiltskin |
| **Poly(A)-binding protein** | | | | | | |
| 38 | AAEL027737 | RRM | 262 | 1 | polyadenylate-binding protein-like | Dmel_CG5119, pAbp |
| 39 | AAEL010318 | 4RRM, PolyA | 628 | 1 | Polyadenylate-binding protein | Dmel_CG5119, pAbp |
| 40 | AAEL020311 | RRM | 249 | 1 | polyadenylate-binding protein 2 | Dmel_CG2163, pAbp2 |
| 41 | AAEL024447 | 2RRM | 433 | 1 | polyadenylate-binding protein-like | Dmel_CG3151 |
| 42 | AAEL007013 | RRM | 175 | 1 | polyadenylate-binding protein 2 | Dmel_CG5119, pAbp |
| **Cytoplasmic polyadenylation element binding proteins (CPEB)** | | | | | | |
| 43 | AAEL019844 | 2RRM | 873 | 2 | cytoplasmic polyadenylation element binding protein 2 | Dmel_CG43782, orb2 |
| 44 | AAEL002422 | 2RRM | 878 | 15 | cytoplasmic polyadenylation element binding protein 1 (CPB) | Dmel_CG10868, oo18 RNA-binding |
| **ELAV like proteins** | | | | | | |
| 45 | AAEL019474 | RRM_3 Bruno, RRM_1 | 688 | 5 | GBP Elav-like family member 2 | Dmel_CG31762, Bruno |
| 46 | AAEL020488 | 2RRM | 391 | 6 | CUGBP GBP Elav-like family member 4 | Dmel_CG43744, Bruno 3 |
| 47 | AAEL006675 | 3RRM | 349 | 5 | ELAV-like protein 3 | Dmel_CG4396, found in neurons |

| | | | | | | |
|---|---|---|---|---|---|---|
| 48 | AAEL010567 | 3RRM | 346 | 1 | ELAV-like protein 2 | Dmel_CG4211, no-on-transient A product form II |
| 49 | AAEL008164 | 3RRM | 363 | 3 | protein elav-like | Dmel_CG4262, embryonic lethal abnormal vision |
| **Eukaryotic translation initiation factor** | | | | | | |
| 50 | AAEL007718 | RRM, 2eIF2A | 688 | 1 | Eukaryotic translation initiation factor 3 subunit B (eIF3b | Dmel_CG4878, eIF3-S9, |
| 51 | AAEL012661 | eIF3g, RRM | 272 | 1 | Eukaryotic translation initiation factor 3 subunit G (eIF3g) | eIF3g1, Dmel_CG8636 |
| 52 | AAEL009646 | RRM | 582 | 2 | eukaryotic translation initiation factor 4B-like | Dmel_CG10837, eIF4B |
| 53 | AAEL004010 | RRM | 333 | 4 | eukaryotic translation initiation factor 4H-like | Dmel_CG4429, eIF4H1 |
| **Nucleolysin** | | | | | | |
| 54 | AAEL009072 | 2RRM | 493 | 3 | nucleolysin TIA-1 | Dmel_CG34354 |
| **Peptidyl-propyl cis-trans isomerase** | | | | | | |
| 55 | AAEL007273 | RRM, Proisomerase | 304 | 1 | peptidyl-prolyl cis-trans isomerase E | Dmel_CG4886, cyclophilin-33 |
| 56 | AAEL005690 | Proisomerase, RRM | 664 | 1 | peptidyl-prolyl cis-trans isomerase sig-7 | Dmel_CG5808 |
| **Musashi like proteins** | | | | | | |

| | | | | | | |
|---|---|---|---|---|---|---|
| 57 | AAEL019841 | 2RRM | 545 | 2 | RNA-binding protein Musashi homolog Rbp6 | Dmel_CG5099, musashi |
| 58 | AAEL000729 | RRM | 333 | 5 | RNA-binding protein Musashi homolog Rbp6 | Dmel_CG32169, Rbp6 |
| **Splicing factors** | | | | | | |
| 59 | AAEL023115 | RRM | 230 | 1 | RNA-binding protein 7-like | Dmel_CG3780, spliceosomal protein on the X |
| 60 | AAEL005046 | 3RRM | 575 | 1 | RNA binding protein 39 | Dmel_CG11266, caper |
| 61 | AAEL007239 | G_patch, RRM | 419 | 1 | splicing factor 45 | Dmel_CG17540, Spf45 |
| 62 | AAEL005909 | RRM | 126 | 1 | splicing factor 3B subunit 6-like protein | Dmel_CG13298, splicing factor 3b subunit 6 |
| 63 | AAEL013795 | 2RRM | 362 | 1 | splicing factor3b subunit 4 | Dmel_CG3780, spliceosomal protein on the X |
| **Transformer proteins** | | | | | | |
| 64 | AAEL006416 | RRM | 273 | 5 | transformer 2 | Dmel_CG43744, bruno 3 |
| 65 | AAEL004293 | RRM | 244 | 1 | transformer 2 | Dmel_CG10128, tra2 |
| 66 | AAEL027148 | RRM | 223 | 2 | transformer-2 protein homolog beta-like | Dmel_CG10466 |
| 67 | AAEL009224 | RRM | 282 | 4 | transformer-2 protein homolog alpha | Dmel_CG10128, tra2 |
| **La proteins** | | | | | | |
| 68 | AAEL007853 | LA, RRM, RRM_3 | 537 | 1 | la-related protein 7 | Dmel_CG42569, La autoantigen-like 2 |
| 69 | AAEL003664 | LA, RRM, RRM_3 | 393 | 1 | la protein homolog | Dmel_CG10922, La autoantigen-like |
| 70 | AAEL015118 | LA, RRM | 1942 | 24 | la-related protein | Dmel_CG11505, La-related protein 4B |

| | | | | | | |
|---|---|---|---|---|---|---|
| **SRA stem-loop interacting RBPs** | | | | | | |
| 71 | AAEL020196 | 6RRM | 1389 | 2 | SRA stem-loop interacting RNA binding protein 2 | Dmel_CG8021, SRA stem-loop interacting RNA binding protein 2 |
| 72 | AAEL021950 | 7RRM | 1393 | 1 | SRA stem-loop interacting RNA binding protein 1 | Dmel_CG33714 |
| 73 | AAEL006645 | RRM | 91 | 1 | SRA stem-loop-interacting RNA-binding protein, mitochondrial | Dmel_CG8021, SRA stem-loop interacting RNA binding protein 2 |
| **Sex lethal** | | | | | | |
| 74 | AAEL011150 | 2RRM | 285 | 2 | sex-lethal | Dmel_CG43770, sex lethal |
| **Other characterised RBPs** | | | | | | |
| 75 | AAEL012618 | 2RRM | 552 | 1 | RNA-binding protein 34 | Dmel_CG4211, no-on-transient A product form II |
| 76 | AAEL026522 | 3RRM | 638 | 1 | RNA-binding protein 28 | Dmel_CG4806 |
| 77 | AAEL006907 | RRM | 198 | 1 | probable RNA-binding protein 46 | Dmel_CG17838, syncrip |
| 78 | AAEL000272 | RRM | 198 | 2 | probable RNA-binding protein 18 | Dmel_CG14414 |
| 79 | AAEL003407 | RRM | 283 | 5 | RNA-binding protein 24-A | Dmel_CG6354, ribonuclear protein at 97D |
| 80 | AAEL006748 | RRM | 165 | 1 | RNA-binding protein 8A | Dmel_CG8781, tsunagi |
| 81 | AAEL004075 | 6RRM | 860 | 1 | RNA binding motif protein | Dmel_CG3335 |
| 82 | AAEL011651 | RRM | 621 | 2 | probable RNA binding protein | Dmel_CG14230 |

| | | | | | | |
|---|---|---|---|---|---|---|
| 83 | AAEL010665 | 4RRM | 491 | 1 | RNA-binding protein 45 | Dmel_CG1316 |
| 84 | AAEL000385 | 4RRM | 412 | 2 | RNA-binding protein 45 | Dmel_CG1316 |
| 85 | AAEL023022 | RRM, PWI | 882 | 1 | RNA binding protein 25 | Dmel_CG4119 |
| 86 | AAEL012171 | 2RRM, SPOC | 756 | 4 | putative RNA-binding protein 15 | Dmel_CG2910, spenito |
| 87 | AAEL010642 | 1RRM | 328 | 3 | RNA binding protein 42 | Dmel_CG2931 |
| 88 | AAEL019795 | 3RRM | 929 | 4 | RNA-binding protein fusilli | Dmel_CG8205, fusilli |
| 89 | AAEL027193 | RRM | 138 | 1 | RNA-binding motif protein, X-linked 2 | Dmel_CG10466 |
| 90 | AAEL012119 | RRM | 239 | 1 | nucleolin | Dmel_CG14414 |
| 91 | AAEL001997 | RRM | 267 | 1 | nucleolin 1 | Dmel_CG3594 |
| 92 | AAEL019934 | RRM | 680 | 1 | mucin-5AC | Dmel_CG10084, Swm |
| 93 | AAEL024670 | RRM | 1900 | 1 | histone-lysine N-methyltransferase SETD1 | Dmel_CG40351, SET domain containing 1 |
| 94 | AAEL028101 | RRM_1 | 244 | 6 | Couch potato | Dmel_CG43738, couch potato |
| 95 | AAEL004415 | 3RRM | 596 | 5 | poly(U)-binding-splicing factor half pint | Dmel_CG12085, Half pint |
| 96 | AAEL019945 | 5HAT, 2RRM, Lsm_interact | 891 | 1 | squamous cell carcinoma antigen recognized by T-cells 3 | Dmel_CG3312, RNA-binding protein 4F |
| 97 | AAEL019944 | 5HAT, 2RRM | 886 | 1 | squamous cell carcinoma antigen recognized by T-cells 3 | Dmel_CG3312, RNA-binding protein 4F |
| 98 | AAEL008938 | RRM | 441 | 1 | polymerase delta-interacting protein 3 | Dmel_CG6961 |
| 99 | AAEL009430 | 4RRM, SPOC | 5680 | 10 | protein split ends | Dmel_CG18497, split ends |

| | | | | | | |
|---|---|---|---|---|---|---|
| 100 | AAEL012100 | RRM | 434 | 1 | MKI67 FHA domain-interacting nucleolar phosphoprotein | Dmel_CG34362, trivet |
| 101 | AAEL017030 | RRM | 193 | 1 | repressor splicing factor 1 (rsf1) | Dmel_CG5655, repressor splicing factor 1 |
| 102 | AAEL011988 | 2RRM | 318 | 1 | tRNA selenocysteine-associated protein (secp43) | Dmel_CG15440, tRNA selenocysteine associated protein |
| 103 | AAEL013869 | RRM, CSTF2_hinge, CTSF_C | 399 | 1 | Cleavage stimulation factor subunit 2 tau variant-like | Dmel_CG7697, cleavage stimulation factor 64 kDa subunit |
| 104 | AAEL017164 | RRM | 631 | 3 | cleavage and polyadenylation specific factor subunit | Dmel_CG7185, cleavage and polyadenylation specific factor 6, isoform |
| 105 | AAEL001352 | SAP, RRM | 811 | 1 | scaffold attachment factor b (SAFB)-like transcription modulator | Dmel_CG6995, scaffold attachment factor B |
| 106 | AAEL004053 | Nup35_RRM_2 | 332 | 1 | nucleoporin NUP53 | Dmel_CG6540, nucleoporin 35kDa Nup35 |
| 107 | AAEL024343 | 2RRM | 530 | 1 | TAR DNA-binding protein 43-like | Dmel_CG10327, TAR-binding protein |
| 108 | AAEL026243 | RRM, FoP_duplication | 261 | 1 | THO complex subunit 4-A | Dmel_CG1101, RNA and export factor binding protein 1 |
| 109 | AAEL013723 | 2RRM_5 1RRM_8 | 620 | 12 | polypyrimidine tract-binding protein 2 | Dmel_CG31000, hephaestus |
| 110 | AAEL014999 | 2RRM | 542 | 15 | protein alan Shepard | Dmel_CG32423, alan shepard |
| 111 | AAEL019836 | RRM | 183 | 8 | protein boule-like | Dmel_CG4760, boule |

| | | | | | | |
|---|---|---|---|---|---|---|
| 112 | AAEL007449 | M_domain, RRM | 1529 | 5 | protein Gawky-like | Dmel_CG31992, gawky |
| 113 | AAEL019609 | UBA, M_domain, RRM | 1460 | 3 | protein Gawky-like | Dmel_CG31992, gawky |
| 114 | AAEL005528 | NTF2, RRM | 744 | 2 | ras GTPase-activating protein-binding protein 1-like | Dmel_CG9412, rasputin |
| 115 | AAEL000674 | RRM, methyltransferase | 625 | 1 | tRNA (uracil-5-)-methyltransferase homolog A | Dmel_CG3808 |
| 116 | AAEL010815 | 2RRM, FCH, muHD | 1345 | 12 | F-BAR domain only protein 2 | Dmel_CG8176 |
| 117 | AAEL025258 | 2RRM | 652 | 2 | HIV Tat-specific factor 1 | Dmel_CG6049 barricade |
| 118 | AAEL017116 | 2RRM | 600 | 5 | hrp65 protein | Dmel_CG4211, no-on-transient A product form II |
| 119 | AAEL012243 | 6RRM | 737 | 1 | uncharacterised | Dmel_CG10328, nonA-like |
| 120 | AAEL006876 | RRM, 4KH | 541 | 5 | igf2 mRNA binding protein 1, putative | Dmel_CG1691, IGF-II mRNA-binding protein |
| 121 | AAEL026633 | RING, RRM | 1368 | 13 | mediator of RNA polymerase II transcription subunit 13 | Dmel_CG31716, CCR4-NOT transcription complex subunit 4 |
| 122 | AAEL003077 | RRM | 279 | 2 | negative elongation factor E | Dmel_CG5994, negative elongation factor E |
| 123 | AAEL006135 | RRM | 159 | 1 | Nuclear cap-binding protein subunit 2 | Dmel_CG12357, cap binding protein 20 |

| | | | | | | |
|---|---|---|---|---|---|---|
| 124 | AAEL014649 | RRM, HGTP_anticodon | 452 | 1 | nuclear receptor coactivator 5-like | Dmel_CG8614, neosin, |
| 125 | AAEL000042 | DnaJ, RRM_1 | 300 | 1 | DnaJ homolog subfamily C member 17-like | Dmel_CG10128, tra2 |
| 126 | AAEL010200 | RRM | 697 | 2 | ecto-NOX disulfide-thiol exchanger 2 | Dmel_CG10948 |
| 127 | AAEL003916 | RRM | 296 | 1 | glycine-rich RNA binding protein GRP2A | Dmel_CG8597, lark |
| **Unknown RBPs** | | | | | | |
| 128 | AAEL022113 | 4RRM | 1229 | 1 | uncharacterized | No |
| 129 | AAEL019864 | 5RRM | 1510 | 2 | uncharacterized | Dmel_CG7879 |
| 130 | AAEL004699 | 4RRM | 1006 | 1 | uncharacterized | No |
| 131 | AAEL022876 | RRM, ZnF_C2H2 | 512 | 7 | uncharacterized | Dmel_CG42458 |
| 132 | AAEL023907 | 5RRM | 1381 | 1 | uncharacterized | Dmel_CG17136, RNA-binding protein 1 |
| 133 | AAEL025927 | RRM | 758 | 1 | uncharacterized | Dmel_CG14937 |
| 134 | AAEL022693 | 6RRM | 1305 | 1 | uncharacterized | Sex lethal, Dmel_CG43770 |
| 135 | AAEL019879 | RRM_1 | 598 | 3 | uncharacterized | Dmel_CG3691, painting of fourth |

**Supplementary figure:**

**Figure S1.** Amino acid sequence alignment of squid protein and its orthologs in *Aedes aegypti*. Jalview has been used for alignment. Identical residues are marked with dots whereas high conservation in the amino acid sequences of both proteins is shown in bright yellow color and higher numerical value.

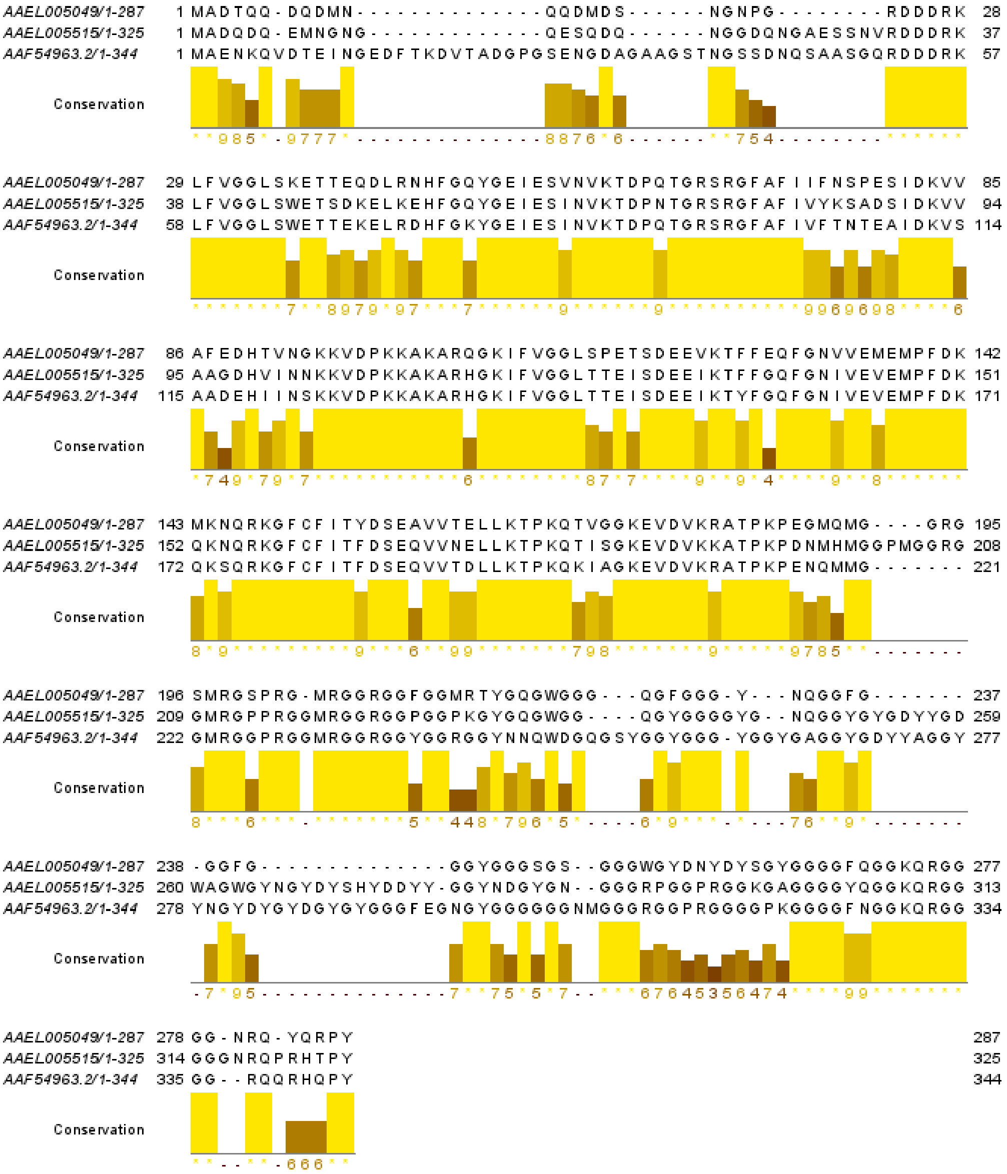
